# Quorum sensing as a mechanism to harness the wisdom of the crowds

**DOI:** 10.1101/2022.03.16.481998

**Authors:** Stefany Moreno-Gámez, Michael E. Hochberg, G. S. van Doorn

## Abstract

Bacteria release and sense small molecules called autoinducers (AIs) in a process known as ‘quorum sensing’ (QS). The prevailing interpretation of QS is that by sensing AI concentrations, bacteria estimate population density to regulate the expression of functions that are only beneficial when carried out by a sufficiently large number of cells. However, a major challenge to this interpretation is that the concentration of AIs strongly depends on the environment, often rendering AI-based estimates of cell density unreliable. Here we propose an alternative interpretation of QS, where bacteria, by releasing and sensing AIs, harness social interactions to sense the environment as a collective. As shown by a model, this functionality can explain the evolution of QS, and results from individuals improving their estimation accuracy by pooling many imperfect estimates – analogous to the ‘wisdom of the crowds’ in decision theory. Importantly, our model reconciles the observed dependence of QS on both population density and the environment and explains why several QS systems regulate the production of private goods.

## Introduction

Quorum sensing is a process whereby bacteria synthesize small molecules known as autoinducers (AIs) that are either passively or actively released into the extracellular space. The same bacteria that produce AIs also sense and respond to the extracellular concentration of AIs via specialized receptors that bind these molecules and initiate signal transduction cascades once the AI concentration is above a certain threshold. These signaling cascades have been well described in many systems and regulate processes such as biofilm formation, virulence, competence and sporulation (1–3). Despite the detailed understanding of the molecular mechanisms underlying various QS systems, the adaptive value and evolutionary origin of QS are less understood (4, 5). The prevailing functional interpretation of QS states that bacteria engage in releasing and sensing AIs in order to monitor population density. This would ensure that individuals express QS-regulated traits only when there is a sufficiently high number of other individuals also expressing them (hence the term ‘quorum’) (2, 3).

This explanation is based on two premises that have been challenged in light of accumulating evidence on the diversity and complexity of QS systems. The first is that the benefit of expressing a QS-regulated trait for an individual increases with population density. There is some evidence of this premise for QS systems that regulate traits involving the production of ‘public goods’ (e.g. extracellular proteases), since the benefit of expressing such traits increases if more cells also express them (6). However, QS also regulates the expression of ‘private’ functions, such as intracellular metabolic enzymes, competence or sporulation (7) that are not shared with other cells and thus provide density-independent benefits (6, 8). To the extent that private functions are more important than public goods to individual fitness, it is unclear why bacteria would regulate such functions by monitoring cell density.

The second premise is that bacteria can reliably estimate population density and the potential for fitness gains by sensing the local AI concentration. This assumption has been notably challenged by studies in different QS systems demonstrating that the relationship between cell density and the concentration of AIs can be contingent on environmental conditions. The best-known environmental factor mediating this relationship is the diffusivity of the extracellular environment. For instance, at sufficiently low diffusivity the AI concentration can result in the quorum for QS induction to be a single cell (9). These and other observations led to the ‘diffusion sensing’ hypothesis, which states that bacteria release AI to test diffusivity and regulate the secretion of costly molecules into the extracellular environment (10). However, given that many other factors such as pH, oxygen and antibiotic stress can affect QS systems (11–13), emphasizing diffusion as the main functional driver of QS likely underestimates the complexity of QS regulation. An alternative, more integrative perspective, acknowledges that multiple biotic and abiotic factors influence QS and responding to a combination of these factors rather than to a single one better explains the functional role of QS for bacteria in nature (4, 13, 14).

If QS is indeed an adaptive mechanism to respond to a combination of abiotic and biotic environmental factors, this raises the question of why bacteria would evolve to employ collective sensing of environmental information over direct individual sensing. Here we propose that bacteria benefit from regulating gene expression through QS because cell-to-cell communication allows individuals to collectively determine the state of the environment by pooling information at spatially relevant scales, which in turn enables them to make more reliable decisions about when to upregulate the expression of QS-controlled traits. According to this hypothesis, cells sense their environment using various mechanisms and encode this information in the rate of AI production - an assumption supported by observations from multiple QS systems (12, 13, 15–19). Then, by secreting AIs and monitoring their extracellular concentrations, cells can share private estimates of environmental conditions and gain access to a ‘pooled’ estimate of the environment – analogous to the “wisdom of crowds” in decision theory, whereby noise in individual estimates of the environment promotes the use of group consensus (20, 21). This hypothesis is not exclusive with the prevailing paradigm that bacteria use QS to coordinate gene expression at high cell densities. Nevertheless, we show here that the benefits derived from collective sensing are sufficient to explain the evolution of QS.

## Model

We study the evolution of QS in fluctuating environments, where the estimates of environmental conditions by individual bacteria are noisy. Our model assumes the simplest possible internal network of feedback regulation (Figure 1a), whereby a gene product *A* promotes its own transcription (Supplementary Information). We parametrize this simple positive feedback network such that there are two possible stable states: an ‘OFF’ state where *A* is expressed at a low basal level, and an ‘ON’ state where *A* is expressed at a much higher level than in the OFF state (Figure S1). This is an approximation of systems where bacteria use QS to modulate all-or-nothing programs of gene regulation; this includes decisions such as sporulating or becoming competent or virulent (22–25). We assume that bacteria can exchange *A* with the extracellular environment by passive diffusion through the cellular membrane, which is consistent with how QS works in many Gram-negative bacteria that do not have dedicated transporters for QS signals (26). Hence, *A* acts both as a product of the gene regulatory network and as a QS signal. Furthermore, bacteria in our model can evolve a parameter *c* that determines membrane permeability to *A* and thus the degree of interaction among cells. In nature, membrane permeability can change through variations in the length or biochemical structure of AIs, as well as by the evolution of active mechanisms for AI secretion or transport (e.g. carrier proteins) (27–29).

**Figure 1.**
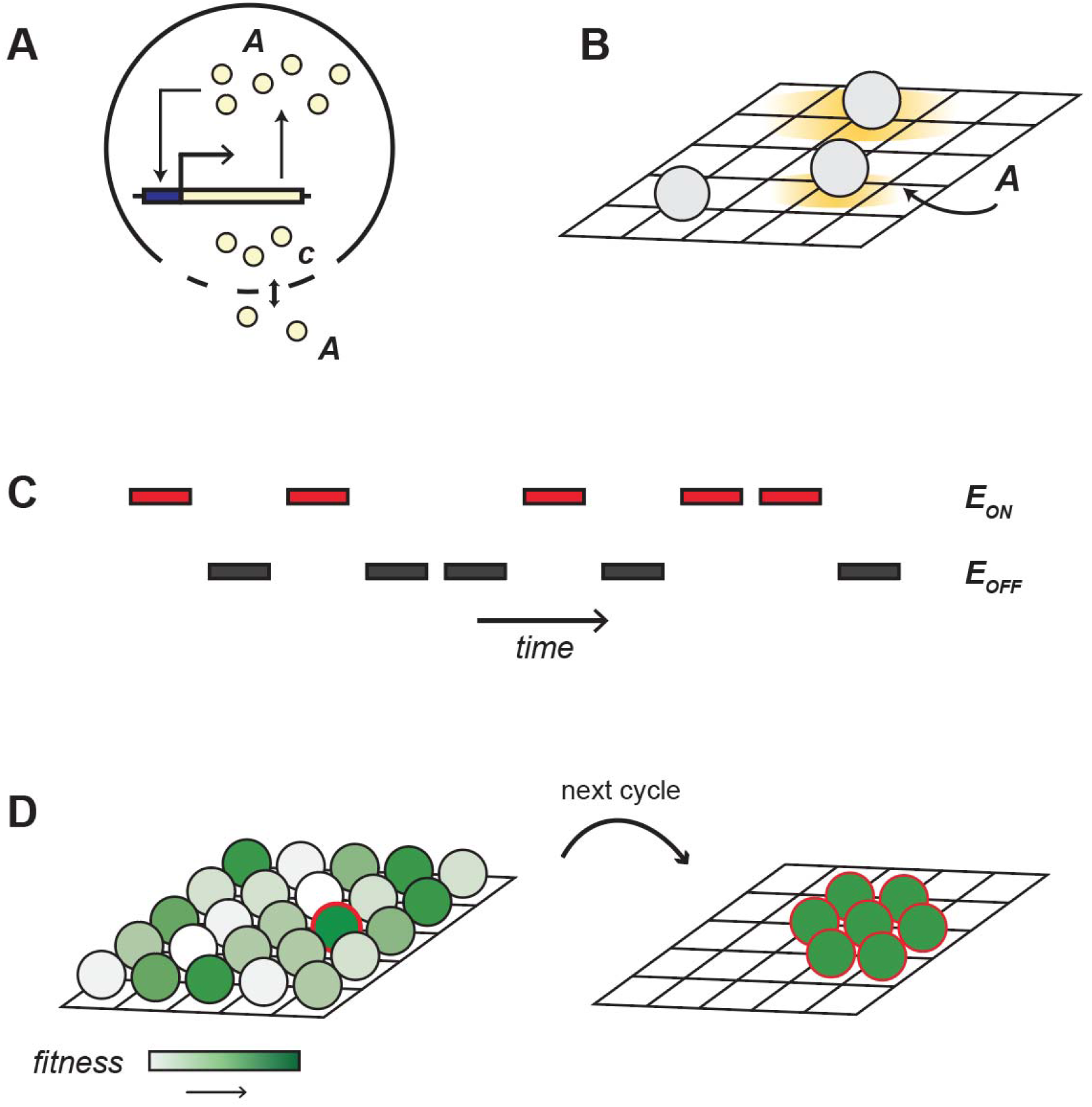
Model structure. (**A**) **Gene regulatory network.** Internally, a bacterial cell has the simplest gene network of positive feedback regulation, where *A* promotes its own transcription. In addition to this regulatory network, bacteria exchange *A* with the extracellular environment by passive diffusion through the membrane at a rate proportional to the evolvable parameter *c.* (**B**) **Diffusion of gene product *A***. At each timestep, the intracellular concentration of *A* i updated for every cell, and the extracellular concentration of *A* is updated according to a diffusion process with diffusion constant *D* over the 2-D grid (bacteria occupy the whole 2-D grid, but only 3 cells are shown here for illustration). The yellow halo shows how *A* leaks from a cell and diffuses over the 2-D grid, and the three cells illustrate different scenarios: (bottom) a cell with *c=0* that does not exchange *A* with the extracellular environment; (center) a cell that either has a low value of *c* or lives in an environment where diffusivity *D* is low; (top) a cell with a high value of *c*, or that lives in an environment with high diffusivity. (**C**) **Environmental fluctuations.** The environment experienced by each cell on the grid fluctuates randomly between two states, *E_ON_* and *E_OFF_*. In generations when the environment is in the *E_ON_* state, bacteria maximize their fitness by expressing *A* at a high level, whereas when the environment is in the *E_OFF_* state, fitness is maximal when *A* is produced at a low basal level. The fitness of every cell is calculated at the end of every environmental cycle as the difference between the level of expression of *A* and its optimal level of expression given the current environmental state, averaged over the duration of the cycle. (**D**) **Fitness values are used to update the grid for the next environmental cycle.** The full grid is repopulated such that individuals with high fitness (fitness increases from white to green) leave more descendants located at or adjacent to their positions on the grid. This is illustrated for a single (red outline) high fitness parent and its offspring. See main text and SI for model details.

Bacteria fully occupy a two-dimensional 50×50 grid over which *A* diffuses with diffusion rate constant *D*. Bacteria evolve through a series of environmental cycles that fluctuate randomly between two equally probable alternative states, *E_OFF_* and *E_ON_* (Figure 1b). In each environmental state there is an optimal level of *A* expression for all individuals in the grid: while *E_OFF_* favors bacteria that do not express *A, E_ON_* favors bacteria with high levels of expression. For instance, *E_ON_* could correspond to an environment where *A* activates an adaptive program to cope with stress (e.g., competence). Activating such a program would not be useful in the absence of the stressor (*E_OFF_*) and thus bacteria would benefit from switching off the production of *A* in this context (Table 1). At the end of each environmental cycle fitness is calculated as the absolute difference between the value of *A* and the current optimal expression level, either *A_ON_* for *E_ON_* or *A_OFF_* for *E_OFF_*, averaged over the cycle duration. Parent cells can bud multiple times during an environmental cycle and the number of offspring produced is determined based on parent fitness relative to other cells in the population. The fitness function has a sigmoidal shape such that individual cells are penalized for errors in determining whether the environment is in the ON or OFF state but not for small numerical deviations from the optimal value of *A* when *c* = 0 (Figure S2 and Supplementary Information).

**Table 1.**
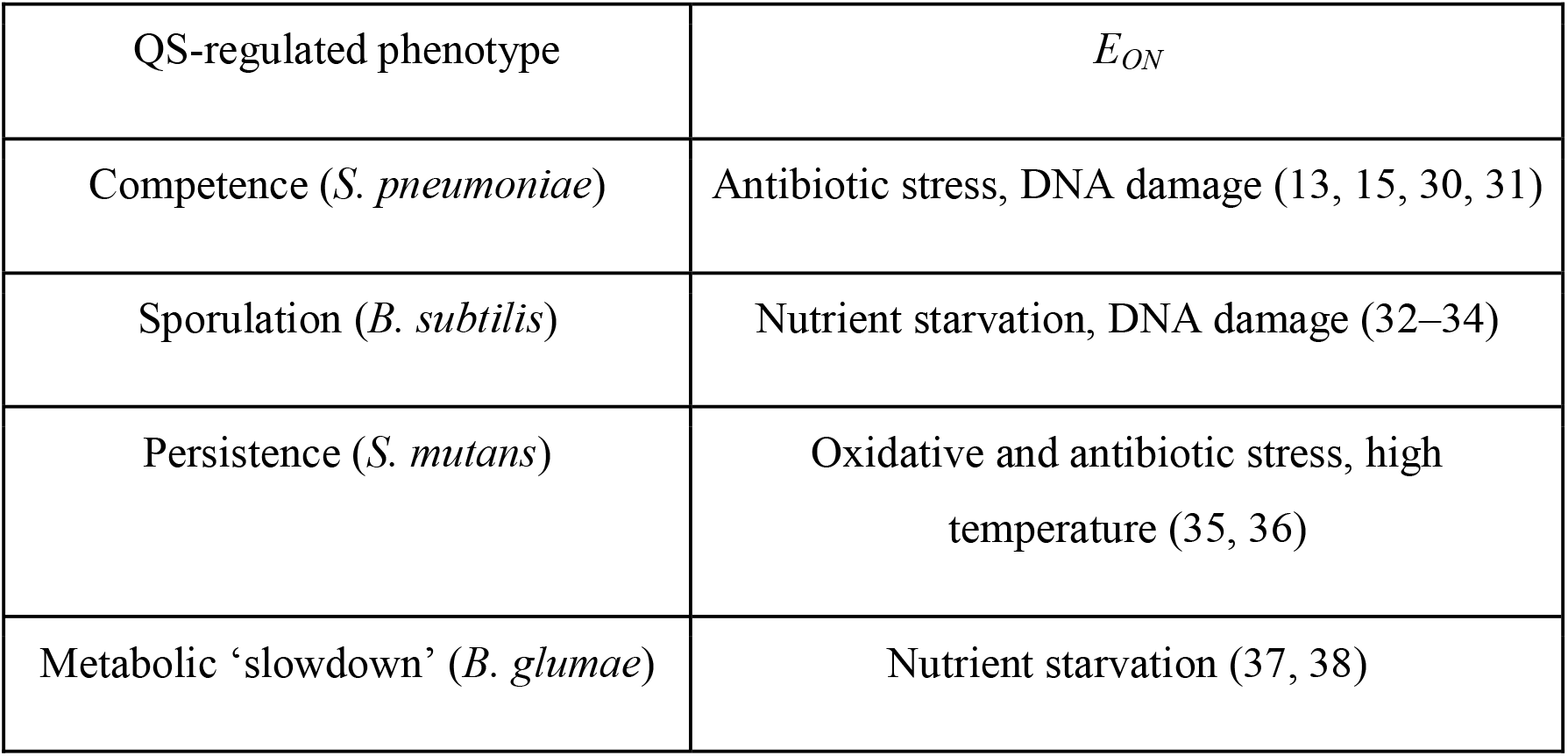
Examples of *E*_*ON*_ environments in different QS systems.

Then, to start a new environmental cycle, the grid is fully repopulated by the daughter cells of the fittest individuals at the end of the previous cycle so generations are non-overlapping. Upon cell birth, *c* mutates with probability *μ*, resulting in *c* increasing or decreasing with equal probability by a fixed step size δ (subject to the constraint that *c* ≥ 0). Finally, daughter cells are placed in the nearest location available to the position of their parent.

Exchanging *A* with the extracellular environment provides cells with information on the initial extracellular concentration of *A,* which could potentially be beneficial if this concentration is informative of the current environmental state. To prevent such benefits (which do not result from cell-cell communication) from biasing the outcome towards the evolution of high *c*, we implement initial conditions for the extracellular concentration of *A* that are uninformative to cells. In particular, we assume that every timestep the initial concentration of *A* is sampled independently for each grid cell from a uniform distribution with mean (*A_OFF_ + A_ON_*) / 2.

Finally, a key assumption of the model is that bacteria can differ in their individual estimates of the environment despite encountering the same environmental regime and having the same internal gene regulation network. Such phenotypic heterogeneity has been documented in several QS systems (e.g., bioluminescence in *Vibrio*, competence in *Bacillus* and virulence in *Listeria*) where actively quorum-sensing isogenic populations contain subpopulations of cells in an OFF state (39, 40). The origin of these phenotypic differences has been partially attributed to stochastic events at the level of expression of quorum-sensing-related molecules, in particular of AIs, response regulators, and proteins involved in the cascades of QS regulation (40–43). In our model, this cell-to-cell variation is captured by assuming that at the start of an environmental cycle each bacterium makes an individual estimate of the state of the environment that is reflected in its internal *A* concentration. We implement this by letting bacteria sample their internal *A* concentration from a (truncated) normal distribution whose mean is the optimal level of *A* expression in the current environment (either *A_ON_* or *A_OFF_*). Sampling from a distribution reflects the assumption that bacterial estimates of the current environment are noisy (to an extent quantified by the variance in the distribution), due to some combination of environmental unpredictability and intrinsic estimation errors.

## Results

Starting from a population where cells do not share information (*c* = 0 initially), we find that *c* increases over time and thus communication readily evolves in the population (Figure 2). Interestingly, although *c* initially increases slowly, there are successive sweeps that lead to a rapid transition towards higher values of *c*, followed again by a slower increase. This occurs because (i) communication becomes beneficial only after a minimum number of neighboring cells are exchanging information (Figure S3) and (ii) once most cells are coupled to each other these benefits increase marginally with the mean value of *c* in the population (Figure S4). Taken together, these results indicate positive frequency-dependent selection on communication and its eventual stabilization.

**Figure 2.**
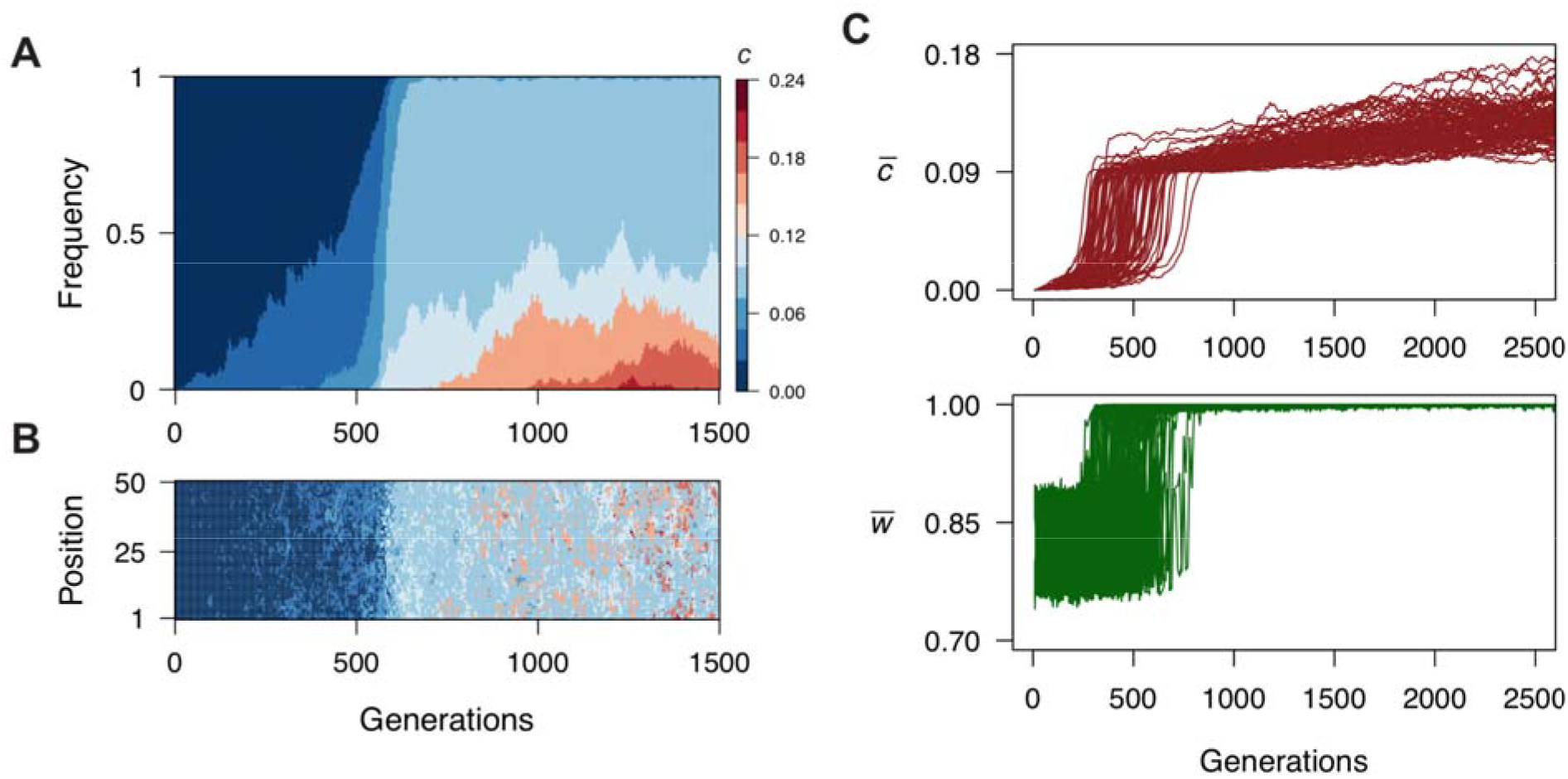
Evolution of QS as a collective sensing mechanism. (**A**) Evolution of th communication parameter *c* across time in a single evolutionary simulation. (**B**) Genetic composition of the bacteria located in a single arbitrary row of the two-dimensional grid tracked through evolutionary time. Although few cells with increased values of *c* emerge in the first 100 generations, they are mostly surrounded by cells with *c*=0 and do not benefit from cell-cell communication. At ~500 generations, the number of cells with c≠0 increases and there are successive sweeps of higher values of *c*. (**C**) Mean *c* (top) and mean population fitness (bottom) across 2500 generations in 100 replicate evolutionary simulations showing that the rapid spread of communication through the population is associated with a rapid increase in the benefit of collective sensing arising once there is a minimum degree of communication in the population. When the mean *c* exceeds 0.09, most cells are communicating to the extent that they can correctly determine the state of the environment and mean fitness approaches 1.0. Thereafter, and due to the sigmoidal shape of the fitness function, evolving higher *c* has a marginal effect on fitness. Parameters: *E_OFF_* = 20, *E_ON_* = 100, *μ* = 0.001, *δ* = 0.03 and *s* = 0.8.

We find that a series of conditions favor the evolution of collective sensing. The first two are related to model assumptions justified previously. First, collective sensing is beneficial only if, on average, cells make an individual estimate sufficiently close to the current state of the environment (Figure S5a). Thus, our model is consistent with a general principle of decision theory known as the Condorcet Jury Theorem. This theorem establishes that for a group of individuals using a majority-rule for decision making, the chance of making the right choic increases with the number of voters only if individuals make the correct choice more often than the incorrect one (44). Second, provided that on average individual estimates of the environment are correct, increased noise in the individual estimates of the environment facilitate the evolution of collective sensing (Figure S5b). In fact, in the extreme scenario where cells could determine the exact state of the environment on their own, there would be no benefit of cell-to-cell communication as a way to improve individual estimates of environmental conditions. Interestingly, the shape of the noise distribution affects the benefit that bacteria derive from communication. In particular, skewed noise distributions where most (but not all) of the cells make an accurate estimate of the environmental state can delay the evolution of cell-to-cell communication (Figure S5c). Finally, intermediate mutational step sizes also favor the evolution of collective sensing because they speed up evolution towards high *c* while keeping the cost of communication low for the first communicators (Figure S6).

In addition to the previous conditions, we find that two features of bacterial interactions facilitate the evolution of collective sensing. First, in the absence of motility, the offspring of a bacterial cell is often located nearby in space. Our model shows that such spatial placement of offspring accelerates the evolution of collective sensing relative to random placement (Figure 3a,c), a result that is consistent with the considerable literature on the importance of space in the evolution of collective behavior (45–47). Collective sensing evolves in our model, because the benefits of cell-to-cell communication are accrued locally through successive environmental generations. Dispersal frustrates this evolution, both in the source habitat of the mother cell and in the target areas to which daughter cells disperse.

**Figure 3.**
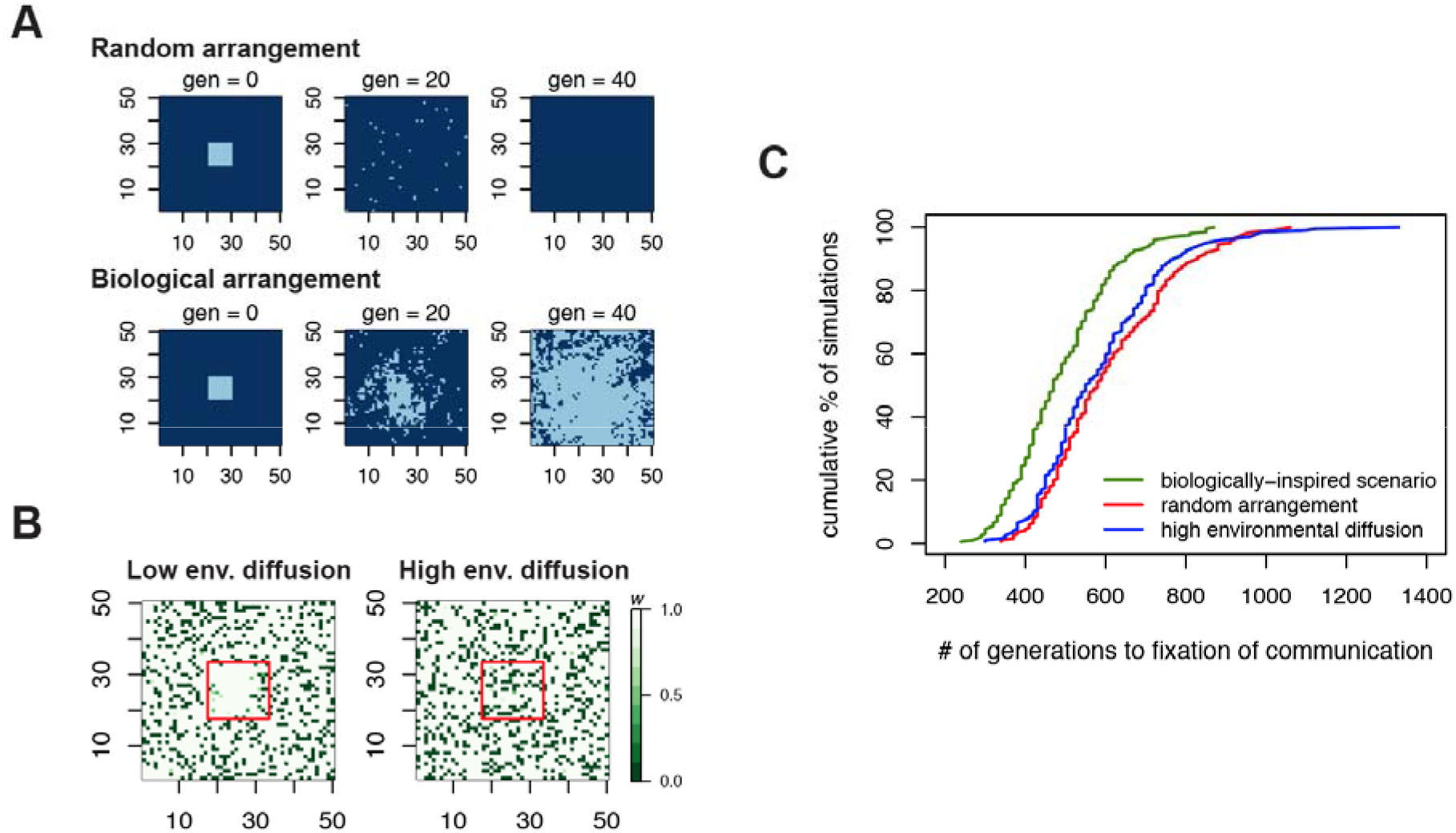
QS, local interactions and the role of environmental diffusion. (**A**) Genetic composition of two populations (shown in the two-dimensional 50×50 grid) that start with a subpopulation of communicators (light blue, c=0.09) surrounded by non-communicators (dark blue, c=0) across 40 generations of selection. In the top populations, the offspring of a cell i placed randomly on the grid, whereas in the bottom populations, offspring occupy a position close to their mother cell, which captures a feature of bacterial reproduction in som environments (See Supplementary Information for description of the algorithm). Random placement of offspring leads to the extinction of communicating cells because bacteria with high *c* only benefit from collective sensing if there are other communicators nearby. (**B**) Individual fitness values in two populations of non-communicators (*c* = 0) that contain a subpopulation of communicators (*c* = 0.09, shown by the red square). When environmental diffusion is low (*D* = 1), the subpopulation of communicators benefits from collective sensing, whereas at high environmental diffusion (*D* = 10), communicating cells are coupled with non-communicators and any fitness benefit of collective sensing disappears. The fitness values are calculated after one generation where the population encounters an *E_ON_* environment. A similar pattern is observed in an *E_OFF_* environment. (**C**) Cumulative distribution of the time to fixation of cell-cell communication in three scenarios: (green) biologically-inspired scenario presented in Figure 2, where the offspring of a cell remain nearby and bacteria interact locally; (red) same scenario as Figure 2 except the offspring of a cell are randomly placed over the spatial grid after reproduction; (blue) same scenario as Figure 2 except the rate *D* of diffusion in the extracellular environment is high (*D* = 10), so bacteria have a long interaction range. 200 simulations are shown per condition and we define communication as fixed in the population when the mean *c* exceeds 0.09 (the first plateau observed in all simulations; Fig 2C). For all panels, unles indicated otherwise, parameters are as in Figure 2. Note that the facilitating effect of the local nature of bacterial interactions on the evolution of collective sensing becomes more evident when the mutational step size is large. When this is the case, collective sensing is strongly impeded if there is high environmental diffusion or if the offspring of a cell is randomly placed over the grid after reproduction, and it does not evolve in any of 100 replicate simulations over 7000 generations.

Second, similar to dispersion of offspring relative to the parent cell, environmental diffusivity also influences the evolution of collective sensing. However, unlike the monotonic negative effect of cell dispersal, extreme high or low diffusion hinders the evolution of cell-to-cell communication (Figure 3). Whereas the absence of diffusion prevents the exchange of information between cells (Figure S7), highly diffusive environments tend to couple communicating bacterial cells with many non-communicators, diminishing the benefit of local assortment and slowing the evolution of communication (Figure 3b,c). Taken together, these results indicate that the evolution of collective sensing is favored when bacterial cells interact locally.

The patterns identified so far occur in spatially homogeneous environments where the only spatial inhomogeneities are in the form of differences in signaling among cells. How might our results be influenced by realistic environmental gradients, similar to those generated by abiotic or biotic processes? To answer this, we studied the role of spatial heterogeneity in the evolution of communication by modeling different degrees of intermixing of the environmental states *E_OFF_* and *E_ON_.* In each generation, the environmental state *E_OFF_* existed in one of two spatial domains with equal probability, and state *E_ON_* existed in the other. These spatial domains were generated using a stochastic spatial pattern generator, where pixels of a grid preferably transition to the state occupied by the majority of their neighbors (Supplementary Information). By running this generator for different numbers of steps starting from an initial random configuration of the grid, we were able to generate either fine or coarse-grained domains that were then used as a basis for simulating environments with high and low heterogeneity, respectively (Figure 4).

**Figure 4.**
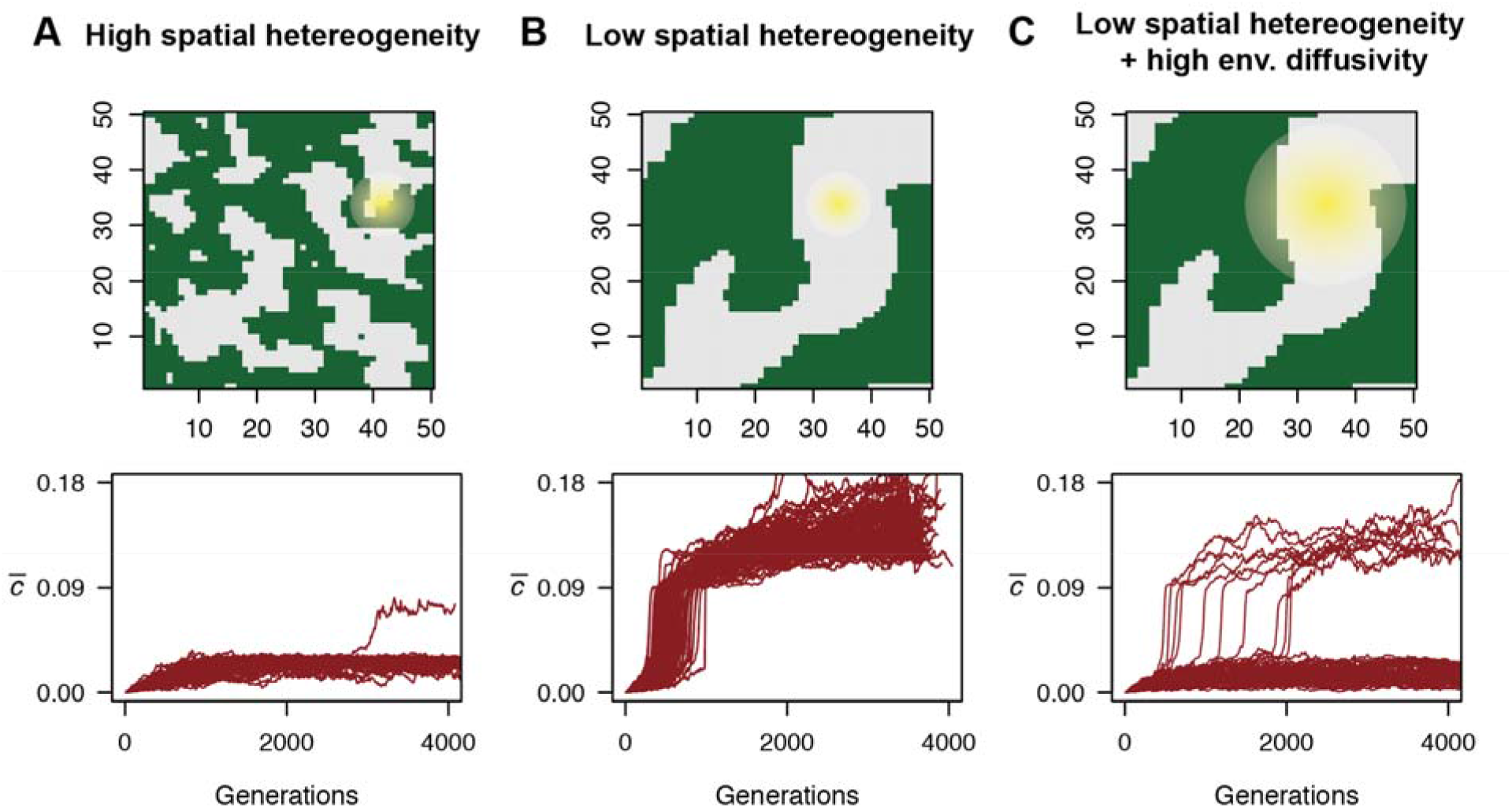
Evolution of QS in spatially heterogeneous environments. (**A**) (Top) Example of the spatial domains (green vs. grey reflecting *E_OFF_* vs. *E_ON_* or vice versa) featuring contrasting environmental states in a single evolutionary simulation with high spatial heterogeneity. The yellow halo represents the neighborhood of interaction of a focal cell; the size of this neighborhood depends on the rate of environmental diffusivity *D*. (bottom) Mean *c* across 4000 generations in 100 replicate evolutionary simulations with high spatial environmental heterogeneity. Cell-to-cell communication evolves only in a very small fraction of the simulations, because communicating cells receive conflicting information from individual experiencing a different environmental state. Panel (**B**) (top and bottom) shows results for a scenario with low environmental heterogeneity. Cell-to-cell communication evolves in all simulations. Panel (**C**) (top) reflects a scenario with low spatial heterogeneity and a high rate of environmental diffusivity, as illustrated by the large size of the yellow halo. Despite coarse spatial heterogeneity, high diffusion increases the coupling between cells experiencing different environmental states, generally undermining the information value of the external autoinducer signal. As a result, cell-to-cell communication evolves in only a small fraction of the simulations (bottom) and takes on average longer to evolve than in a spatially homogeneous environment.

When environmental structure is fine-grained, individuals are very likely to interact with neighbors experiencing different environmental regimes. As a result, they are exposed to misleading information about the state of the environment, which in turn hinders the evolution of collective sensing (Figure 4a). When there is coarse-grained environmental structure, this effect also occurs at domain boundaries. However, cell-cell communication is still beneficial for cells in the center of the spatial domains since they interact with other individuals experiencing the same environmental conditions. Therefore, in contrast to environments with fine-grained structure, collective sensing evolves more readily when spatial heterogeneity is low (Figure 4b). Importantly, these findings are contingent on the size of the interaction neighborhood (yellow halo, Figure 4), which is set by the rate of environmental diffusivity *D*. We illustrate this idea by showing that in the same regime with low levels of spatial heterogeneity, high environmental diffusion can prevent the evolution of communication by increasing the interaction neighborhood of cells (Figure 4c). High diffusion makes it more likely that any given cell is communicating with others experiencing a different environmental state, eroding the information contained in the external concentration of *A*. Thus, when environments vary spatially, the evolution of collective sensing is also favored if bacteria interact at a local scale.

## Discussion

Our model shows that QS can evolve solely as a result of its collective sensing functionality, without the need of a benefit resulting from the coordinated action among cells. This alternative interpretation of QS does not preclude such collective responsive functionality but rather complements the classical view by explaining how QS can be stable in the presence of ‘cheaters’ that do not engage in the action carried by the entire population. One possible explanation is that QS does not only control public functions but also private functions (i.e. functions that offer fitness benefits irrespective of the behavior of neighboring cells). The latter are common in many QS systems (6–8) and a collective sensing interpretation of QS could help explain why these functions are controlled by QS and, in turn, how QS remains protected to some extent from ‘cheaters’.

Analogous to classic collective action, collective sensing is prone to cheating, should sharing information be costly and to any extent independent of receiving information from other cells (48, 49). The latter condition does not hold in our model, where sharing and receiving information are completely coupled because of our choice of modeling *A* as a signal that diffuses passively across the membrane. However, in QS systems in nature, AIs can also be secreted and sensed using dedicated transporters as it is the case for Gram-positive bacteria (50, 51). Thus, in some QS systems, defectors could emerge and reap the benefits of communication without contributing to the public signal, especially if the production of QS signals is costly.

Importantly, our model shows that collective sensing can explain the evolution of QS, because of two essential features of bacterial interaction networks. First, when bacteria divide, their offspring generally remain close in space. This feature not only protects QS from cheater invasion via a kin selection mechanism (52, 53), but as shown here it can facilitate the emergence of a sufficiently large cluster of communicators for collective sensing to be profitable. Second, bacterial interactions occur over short spatial ranges on the order of microns for certain QS systems as reported recently (54–56). We show here that this feature of bacterial interactions could have favored the evolution of collective sensing because emergent communicators (i) are often coupled to their communicating offspring (Figure 3) and (ii) avoid long-range interactions with cells located in different microenvironments that could share deceiving information (Figure 4).

Collective sensing has been proposed as a mechanism for decision-making in other systems, resulting from social interactions among individuals with simple behavioral rules (20, 57). A notable instance in several animal groups is “the wisdom of crowds”, where individuals can improve estimation accuracy by aggregating their separate estimates of the environmental state. Examples of this phenomenon range from nest-site choices by ant colonies (58, 59), to foraging decisions by fish schools (60) and even medical diagnostics (61, 62). Although bacteria do not possess the complex sensing, cognition and feedback mechanisms found in social animals and humans, our work shows that the same collective functionality can arise in populations of bacteria with simple gene regulation networks and could have driven the evolution of quorum sensing as one of the most widely used communication systems in bacteria.

## Acknowledgments

We thank Martin Ackermann, Franz J. Weissing and Simon van Vliet for comments on earlier versions of this manuscript. SMG and GSvD were supported by a Starting Independent Researcher Grant (309555) of the European Research Council and a Vidi fellowship (864.11.012) of the Netherlands Organization for Scientific Research.

