## Supplementary Information for "Quorum sensing as a mechanism to harness the wisdom of the crowds"

#### Contents

**1 Supplementary text** **2**

**2 Supplementary Figures** **5**

### 1 Supplementary text

We study the evolution of cell-to-cell communication in a bacterial population encountering varying environments. The phenotype of a cell is determined by a simple network of positive regulation where a protein $A$ promotes its own transcription (Fig. S1). We model this positive feedback by assuming that the transcriptional regulation of $A$ follows standard Hill kinetics. Bacterial cells inhabit a two-dimensional grid of size $N \times N$ where they can communicate with other cells by exchanging $A$ with the extracellular space. $A$ is exchanged by passive diffusion with a diffusion constant $c$. Based on these assumptions the system of equations describing the intracellular concentration of $A$ and extracellular concentration, $A_E$, is,

$$\frac{d\,A}{d\,t} = \sigma \frac{k_0 + k\,(A/\,K_d)^n}{1 + (A/K_d)^n} + c\,(A_E - A) - d\,A \tag{1}$$

$$\frac{d\,A_E}{d\,t} = c\,(A - A_E)\; + D\,\nabla^2\,A_E \tag{2}$$

where $\sigma$ is the number of proteins produced per transcript of mRNA, $k_0$ is the basal transcription rate when the promoter is not bound to any molecule of $A$, $K_d$ is the dissociation constant, $k$ is the maximal transcription rate, $n$ is the degree of cooperative binding, $d$ is the rate of degradation of $A$ and $D$ is the rate of environmental diffusion. Since most QS systems exhibit bistability we choose the parameter values in a way that there are two stable states at a high and low concentration of $A$ when $c = 0$ (Fig. S1). The parameter values we chose are $\sigma = 10$, $n = 6.75$, $K_d = 60.8$, $d = 0.1$, $k_0 = 0.2$ and $k = 1.03$. We use this set of parameters in all the simulations. Every generation we solve the previous system of equations for a fixed number of time steps $T$ and calculate fitness at the end to determine which individuals will leave offspring in the next generation. We assume periodic boundary conditions for the extracellular diffusion of $A$.

#### Fitness calculation and reproduction

In every generation a bacterial population faces one of two possible environments with equal probability. Each environment has an optimal expression level of $A$, denoted by $A_{\text{OFF}}$ or $A_{\text{ON}}$. $A_{\text{OFF}}$ and $A_{\text{ON}}$ are set at the stable equilibria of the bistable system when $c = 0$ (Fig. S1). At the start of a generation all cells sample their initial intracellular value of $A$ from a truncated normal distribution with mean either $A_{\text{OFF}}$ or $A_{\text{ON}}$ depending on the environment and standard deviation $\sigma_{\text{OFF}}$ or $\sigma_{\text{ON}}$ .The fitness of a cell is determined by how well its intracellular $A$ concentration matches the state of the environment throughout the duration of a generation. The fitness function is,

$$w(\Delta_E) = \frac{1}{1 + e^{s\,(\Delta_E - x)}}$$

where $s$ determines the strength of selection, $\Delta_E = \frac{1}{T}\sum_{t=1}^{T} A_t - A_{\text{ON}}$ (*i.e.* the average difference over the $T$ time steps between $A$ and the optimal level of $A$ expression in the current environmental state, in this example $E_{\text{ON}}$) and $x$ is the the midpoint of the sigmoid curve. For all simulations we set $s = 0.8$ and $x = 25$. For this choice of parameters $w$ has a sigmoidal shape that strongly penalizes cells that are in the incorrect phenotypic state but not cells that slightly deviate from the optimal expression levels (Fig. S2).

At the end of a generation the fitness of every cell is calculated and fitness values are normalized by the total fitness of the population. Then, $N \times N$ cells are randomly sampled using the normalized fitness values to populate the new grid. The algorithm for creating and placing the offspring of a cell in the new grid is the following,

1. Draw a random number to determine if $c$ mutates. If $c$ mutates, draw an additional random number to determine whether the new value of $c$ is $c + \delta$ or $c - \delta$. If $c - \delta < 0$, $c$ does not mutate.

2. Calculate distance of the mother cell to all other cells in the grid. We use a euclidean distance metric.

3. Place the new cell in the closest grid cell to the mother cell that is still empty.

This algorithm is applied to the $N \times N$ vector containing the coordinates of the cells that will reproduce and it ensures that the offspring of a cell remains close to the location of its mother cell. In simulations where the offspring of a cell is randomly placed on the two dimensional grid, the grid is filled by rows in the order that cells appear in the $N \times N$ vector.

#### Spatial heterogeneity

We model variations in the environmental conditions on space by using an Ising model [1] to establish the initial configuration of the environment. Using this model we can vary the scale of spatial heterogeneity from a random configuration to a homogeneous grid. In two dimensions, this model consists of a grid where cells can be in two possible states (-1 or +1). The total energy of the system is determined by whether neighboring cells are in the same or in different state and is given by,

$$H = -J \sum_{<ij>} s_i s_j$$

where $< ij >$ denotes all the pairs of neighboring cells, $s_i$ is the state of the grid cell $i$ and $J$ determines the sign of the interaction. We assume that $J > 0$ so over time the system converges from a random configuration to a configuration where all the cells have the same state.

Starting from a random configuration where each grid cell is assigned to any of the two states with equal probability, we simulated this model using a Metropolis Monte Carlo algorithm for $I$ number of iterations [2]. Briefly, each iteration a grid cell is selected at random and its state is flipped. If the energy of the new configuration is lower than the energy of the old configuration the change in state is accepted and the state of the cell is flipped. If $\Delta E \not< 0$, the state of the cell can still be flipped with probability $e^{-\Delta E/S_T}$, where $S_T$ is a scaling constant. In every iteration, this is repeated $N \times N$ times. Given that the grid configuration will reach equilibrium when all grid cells are in the same state, we can vary the scale of environmental heterogeneity by modifying $I$.

For each simulation run we first determine the configuration of the $N \times N$ grid by running the previous algorithm for $I$ iterations. We use the resulting grid configuration made of the two states, -1 and +1, to determine the state of the environment in each grid cell every generation. At the start of every generation a random number is drawn to assign the environmental states to the grid states. $E_{\text{OFF}}$ and $E_{\text{ON}}$ are assigned to grid cells -1 and +1 or vice versa with equal probability every generation. Each bacterial cell samples an initial intracellular $A$ concentration from a distribution whose mean is determined by the environmental state in the grid cell inhabited by the bacterium.

### 2 Supplementary Figures

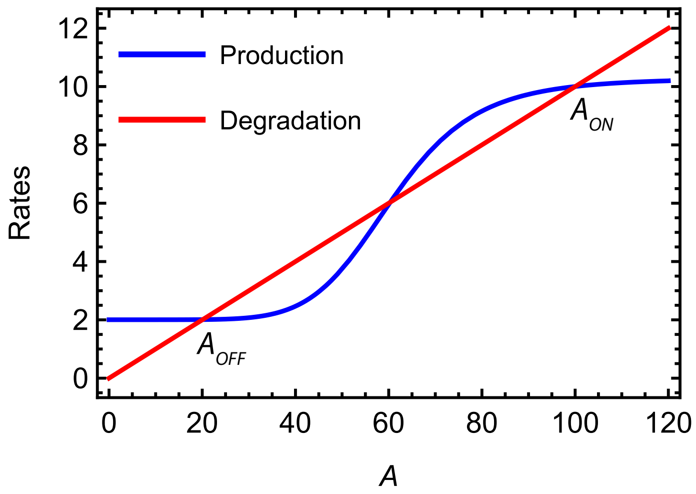


Figure S1: **Steady-states of the model with $c = 0$.** In the absence of diffusion of $A$ to the extracellular space, we chose a set of parameter values that renders the system described by Eq. (1) bistable. The rate of production of $A$ is given by $\sigma \frac{k_0 + k\,(A/\,K_d)^n}{1+(A/K_d)^n}$ and the rate of degradation is $dA$.

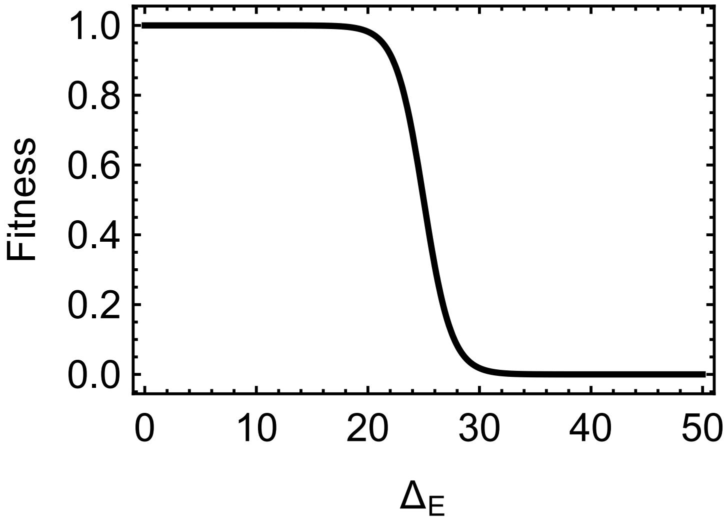


Figure S2: **Fitness function** $w$**.** Function applied at the end of one generation to determine the fitness of a cell depending on the difference $\Delta_E$ between its average value of $A$ and the optimal expression level for the current environmental state.

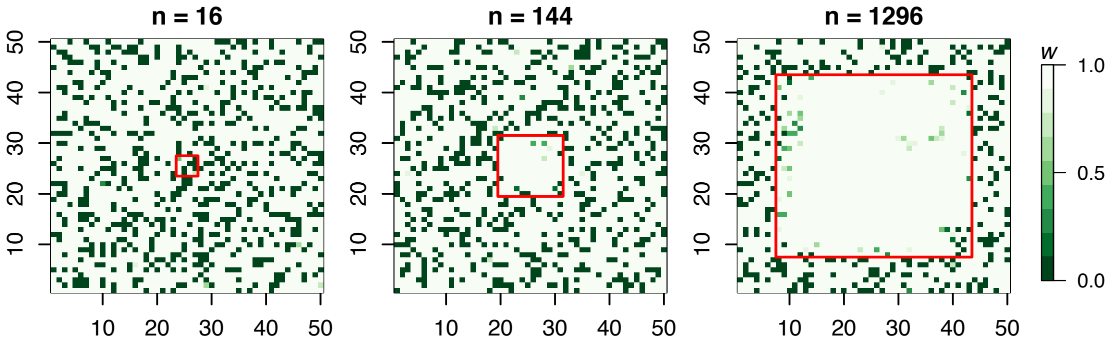


Figure S3: **A cluster of cells communicating is necessary for collective sensing to evolve.** Individual fitness values in three populations of non-communicators that contain a subpopulation of communicating cells. In each panel, the size of this subpopulation is indicated by $n$ and its location in the two-dimensional grid is shown with a red square. Cells benefit from collective sensing once there is a minimum number of communicators. For non-communicators, $c = 0$ and for communicators, $c = 0.09$. Fitness values are calculated after one generation where the population encounters an $E_{\mathrm{ON}}$ environment. A similar pattern is observed in an $E_{\mathrm{OFF}}$ environment. Other parameters are the same as in Fig. 2.

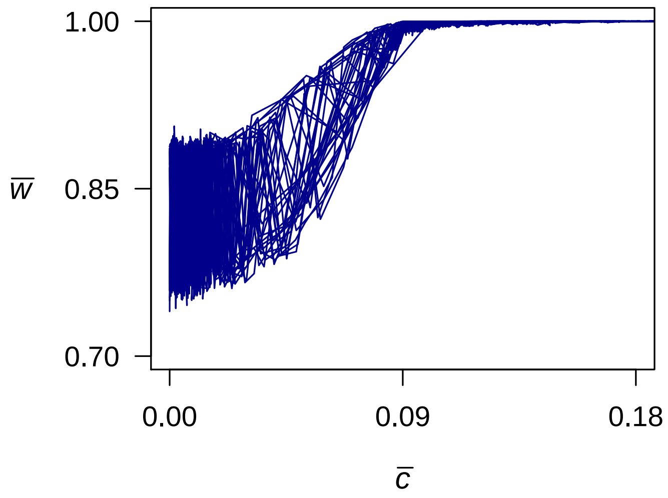


Figure S4: **Fitness increases marginally past a threshold value of $c$.** Mean fitness as a function of the average $c$ for a subset of the evolutionary trajectories presented in Fig. 2. Beyond $\bar{c} = 0.09$ the population is already communicating to the extent that most individuals can correctly estimate the state of the environment every generation. Since the fitness function has a sigmoidal shape that strongly penalizes individuals for making a wrong estimate of the environmental state but not for small deviations from either $A_{\text{OFF}}$ or $A_{\text{ON}}$ (Figure S2), fitness increases marginally with $c$ for $c > 0.09$.

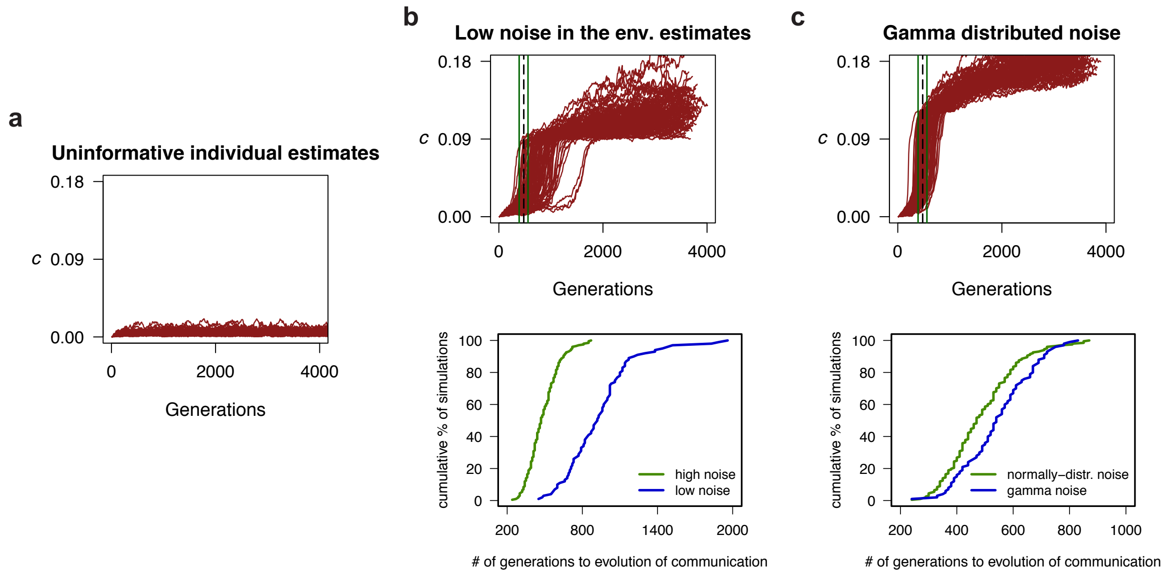


Figure S5: **The evolution of collective sensing depends on the level and structure of environmental noise.** a) When bacteria are unable to individually estimate the state of the environment, collective sensing is not profitable and does not evolve. We model this scenario by assuming that bacteria sample their initial intracellular concentration of $A$ from the same distribution regardless of the state of the environment. b) (*top*) Mean $c$ across 3500 generations in 200 replicate evolutionary simulations with the same parameters as in Fig. 2 but with lower noise in the individual estimates of the environmental conditions ($\sigma_{\mathrm{OFF}} = 10$ and $\sigma_{\mathrm{ON}} = 30$). The mean time for evolution of communication with higher noise in the individual estimates of the environment is shown by the dotted line (Fig. 2, $\sigma_{\mathrm{OFF}} = 30$ and $\sigma_{\mathrm{ON}} = 80$), with the first and third quartiles indicated by green lines. (*bottom*) Cumulative distributions of the time to fixation of cell-cell communication in the two scenarios compared in the figure above. We define communication as fixed in the population when the mean $c$ surpasses 0.09. Since the benefit of collective sensing comes from the error that cells make when estimating environmental conditions, lower noise in such estimates makes communication less profitable and collective sensing takes longer to evolve. c) (*top*) Mean $c$ across 3500 generations in 200 replicate evolutionary simulations with the same parameters as in Fig. 2 when noise in the individual estimates of the environment has the same mean and standard deviation as in Fig. 2 but is gamma-distributed (as opposed to normally-distributed like in Fig. 2). Vertical lines indicate the same as in (b). (*bottom*) Cumulative distributions of the time to fixation of cell-cell communication in the two scenarios compared in the figure above. As in (b), we define communication as fixed in the population when the mean $c$ surpasses 0.09. When noise in the individual estimates of the environment is gamma-distributed some cells can make very inaccurate estimates of the environmental state. When there is communication, these cells can deceive other cells hindering the evolution of collective sensing.

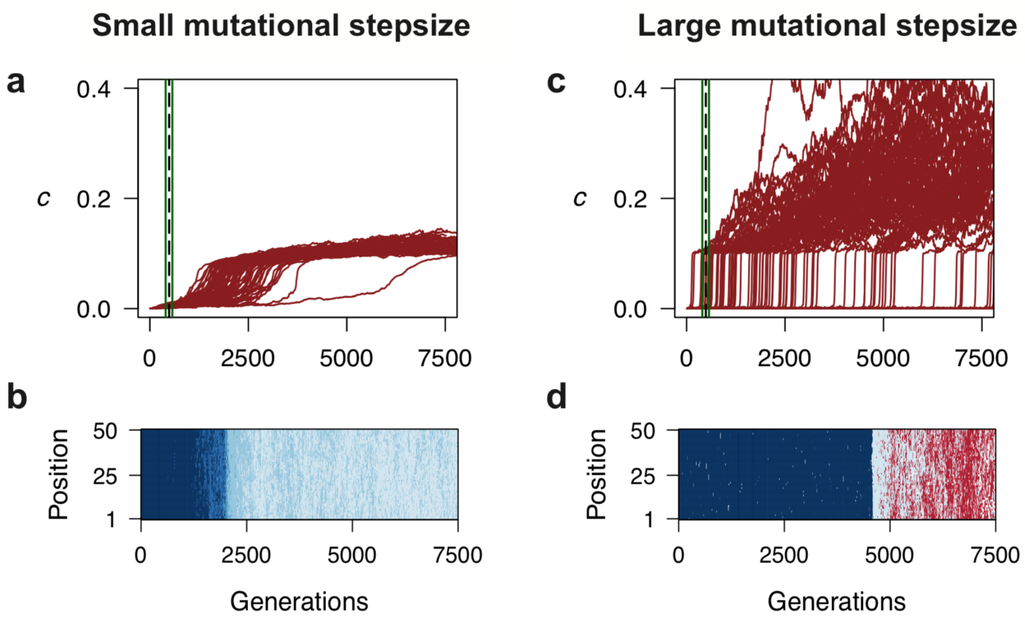


Figure S6: **Intermediate mutational stepsizes favor the evolution of collective sensing.** a) Mean $c$ in 100 replicate evolutionary simulations with the same parameters as in Fig. 2 but a smaller mutational stepsize ($\delta = 0.01$). The mean time for evolution of communication in Fig. 2 ($\delta = 0.03$) is shown by the dotted line with the first and third quartiles shown by green lines. b) Genetic composition of a single row of the two dimensional grid through evolutionary time in one simulation with small mutational stepsize. This illustrates that with a small mutational step it takes very long for a minimum number of communicators with high enough $c$ to emerge relative to an intermediate mutational step (c.f. Fig. 2), which in turn slows down the emergence of collective sensing. c) Mean $c$ in 100 replicate evolutionary simulations with the same parameters as in Fig. 2 but a larger mutational stepsize ($\delta = 0.1$). The solid and dotted lines indicate the same as in panel a). d) Genetic composition of a single row of the two dimensional grid through evolutionary time in one simulation with large mutational step. This illustrates that with a large mutational step communicators arise but go extinct often relative to an intermediate mutational step (like the one in Fig. 2) because in the absence of other communicators high values of $c$ are detrimental since they turn the dynamical system monostable and sensitive to the extracellular concentration of $A$ which is not informative.

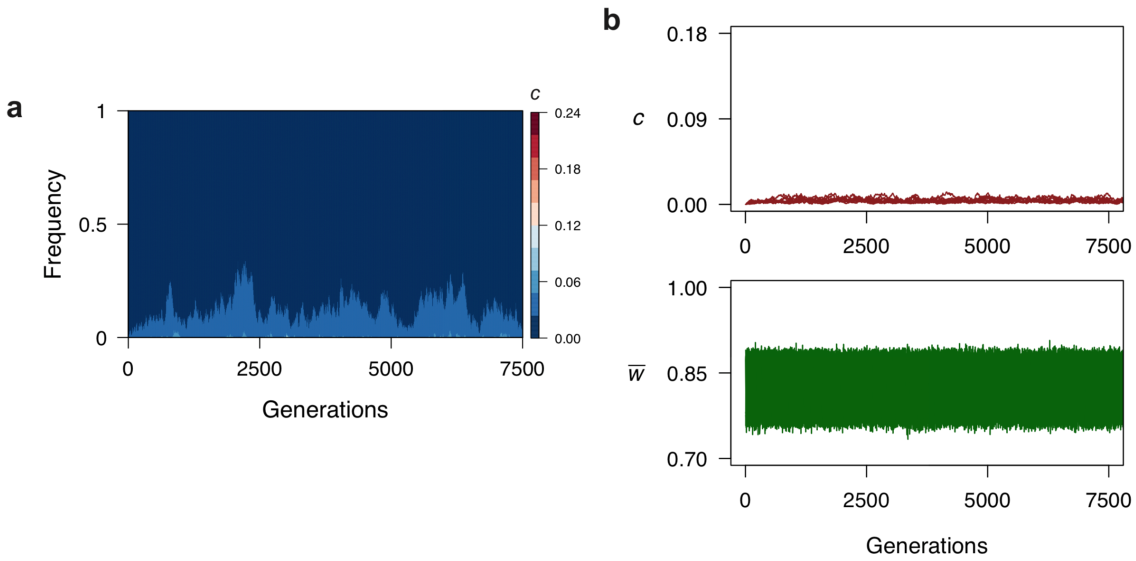


Figure S7: **Communication does not evolve in the absence of extracellular diffusion.** a) Evolution of the communication parameter $c$ across time in a single evolutionary simulation where $D = 0$. b) Mean extracellular diffusion $c$ (top) and mean population fitness (bottom) across 7500 generations in 50 replicate evolutionary simulations. Both panels show that $c$ does not evolve to high values in the absence of environmental diffusion. This shows that cells only benefit from exchanging $A$ with the extracellular environment because they can communicate with other cells and not because they can gather more information on the current environment from the initial extracellular concentration of $A$. All parameters are the same that in Fig. 2 except from $D$.

[2] Michael Adler. *Monte Carlo methods applied to the Ising model*. Diplomica Verlag, Norderstedt, Germany, 2010.
